# Repeated exposure to chlorpyrifos is associated with a dose-dependent chronic neurobehavioral deficit in adult rats

**DOI:** 10.1101/2021.10.28.466295

**Authors:** Ana C. R. Ribeiro, Elisa Hawkins, Fay M. Jahr, Joseph L. McClay, Laxmikant S. Deshpande

## Abstract

Organophosphate (OP) chemicals include commonly used pesticides and also chemical warfare agents, and mechanistically they are potent inhibitors of the cholinesterase (ChE) enzyme. While a chronic low-dose OP exposure does not produce acute cholinergic crises, epidemiological studies report long-term neuropsychiatric issues including depression and cognitive impairments in OP-exposed individuals. Chlorpyrifos (CPF) is one of the most widely used pesticides worldwide. Multiple laboratory studies have reported on either the long-term behavioral effect of a single, high-dose CPF or studied sub-chronic behavioral effects particularly the motor and cognitive effects of repeated low-dose CPF exposure. However, studies on chronic mood and depression-related morbidities following repeated sub-threshold CPF doses that would mimic occupationally-relevant OP exposures are lacking. Here, adult male rats were injected with CPF (1, 3, 5, or 10 mg/kg/d, s.c.) for 21-days. Dependent on the CPF dose, ChE activity was inhibited approximately 60-80% in the blood and about 20-50% in the hippocampus at 2-days after the end of CPF exposures. Following an 11-week washout period, CPF-treated rats exhibited a dose-dependent increase in signs of anhedonia (sucrose preference test), anxiety (open-field and elevated plus-maze), and despair (forced swim test) despite a complete recovery of ChE activity at this stage. We speculate that both cholinergic and non-cholinergic mechanisms could play a role in the development of chronic OP-related depressive outcomes. The proposed CPF exposure paradigm could provide an ideal model to further study molecular mechanisms underlying cause and effect relationships between environmental OP exposures and the development of chronic behavioral deficits.

## Introduction

Depression is one of the most prevalent psychiatric disorders around the world [1]. The etiology of depression is complex and multifactorial, being the cumulative contribution of many genetic and environmental risk factors [2-4]. Recently, exposures to environmental factors such as air pollution, noise, trace pharmaceuticals, and others have received considerable recognition in the literature (reviewed in: [5, 6]). One such environmental factor that has been linked to depression-related outcomes is exposure to pesticides [5-7]. Among the various chemical categories of pesticides, organophosphate (OP) pesticides represent a major class that is used globally. Epidemiological studies have suggested an elevated risk for psychiatric disturbances and suicidal ideations in populations that are exposed to pesticides either occupationally, intentionally, or because of their living conditions [7]. For example, farmworkers [8], sheepherders [9], aircraft personnel [10], or rural residents living in the vicinity of farms [11] are repeatedly exposed to OP-based compounds and report greater incidences of depression. Furthermore, about one-third of the First Gulf War veterans exhibit a chronic disorder known as Gulf War Illness that chiefly includes symptoms of mood disorder, depression, and memory problems [12-14]. Deployment-related exposures to a cocktail of chemicals and pharmaceuticals including repeated exposure to low-level OP pesticides, pyridostigmine tablets, and accidental exposure to the OP nerve gas sarin are thought to contribute to the development of this syndrome [12-14]. In addition to their utility as potent pesticides, OP compounds are also widely used in fuel additives, industrial solvents, pharmaceuticals, and in warfare as lethal nerve agents. Thus, OP exposures are not uncommon and research is needed to understand how they affect biological systems to enhance the risk for depression and related neurologic conditions [15].

Amongst the OP pesticides, chlorpyrifos (CPF) is one of the most widely used worldwide [16]. CPF, like other OP pesticides, produces its effects by blocking the cholinesterase enzymes (ChE) [17]. Within the central nervous system (CNS), this blockade leads to the accumulation of the neurotransmitter ACh at the synapses. The effect of CPF on ChE inhibition is dose-dependent. Thus, within the CNS, high doses of CPF lead to rapid AChE inhibition and produce acute hypercholinergic effects such as salivation, lacrimation, loss of muscle, and sphincter tone. At very high doses, CPF can produce unremitting seizures that can culminate in respiratory depression and rapid death [17]. Survivors of lethal OP exposures often suffer from multiple co-morbid conditions including depression and cognitive dysfunction [18]. On the other hand, sustained low-level OP exposures are not associated with symptoms of acute “cholinergic crises” but epidemiological and clinical studies have shown that repeated, low-dose exposure to OP including CPF across the life span is associated with an enhanced risk for psychiatric disorders. Thus, CPF exposures during gestation [19-22], adolescence [23-25], and adulthood [8, 26, 27] are reported to affect brain morphology [28] and elevate the risk for Attention-Deficit Hyperactivity Disorder (ADHD), autism traits, cognitive disorders and depression [29-32]. Similar findings have been reproduced in laboratory rodent studies. Thus, gestational CPF exposures were reported to produce anxiogenic behavior [33], ADHD-like behavior [34], and exhibit sensorimotor deficits in offspring [35]. Several laboratory studies have also reported the behavioral effects of CPF exposures in young and adult rodents. For example, CPF (1-10 mg/kg/d for 4 weeks) in adult rats produced cognitive impairment and behavioral abnormalities when tested 24-h after the last CPF exposure [36]. Another study reported that CPF (2.5-20 mg/kg from postnatal days 27 to 36) produced depression-like behavioral signs when assessed one week following the last CPF dose [37]. Cognitive deficits along with anxiety traits were noted on days 10 and 14 of a two-week CPF (14.9 mg/kg) exposure paradigm [38]. Another study on behavioral toxicity of repeated CPF exposures showed that CPF (18 mg/kg daily for 14 days or every other day for 30 days) leads to increased impulsivity and reductions in attention when assessed during and up to 30 days post exposure [39, 40].

In summary, multiple laboratory studies have reported on 1) long-term behavioral effects of a single, very high dose CPF exposure, 2) near-term behavioral effects of repeated low-dose CPF exposure or 3) sub-chronic behavioral effects particularly the cognitive effects of repeated low-dose CPF exposure. However, no studies have yet looked at *long-term* mood and depression-related behavioral effects of repeated sub-threshold CPF in adult rats that would mimic occupationally relevant exposures. This is our focus in the current study. We used a battery of rodent behavioral assays that are predictive of signs of mood, anxiety, and despair that are commonly seen in the depression-like condition. Our results indicate that repeated low-dose exposure to OP pesticide CPF at concentrations that did not produce acute hypercholinergic signs or long-term ChE inhibition was associated with signs of chronic behavioral depression.

## Materials and Methods

### Animals

All animal use procedures were in strict accordance with the National Institute of Health (NIH) Guide for the Care and Use of Laboratory Animals. The experimental protocol was approved by Virginia Commonwealth University’s (VCU) Institutional Animal Care and Use Committee (IACUC). Male Sprague-Dawley rats (Envigo, Indianapolis, IN) were ordered at 8-weeks of age and allowed to get acclimated to the housing environment for one week. Animals were housed one per cage at 20-22° C with a 12-hour light-dark cycle (lights on 0600-01800 h), provided with appropriate environmental enrichment (Nyla’s bones), and given free access to food and water. Rats were randomized into five groups of 10 rats each corresponding to various levels of CPF exposures along with its vehicle as described below.

### CPF exposure

CPF (CAS no. 2921-88-2) was obtained from Chem Service, Inc. (Chester, PA) as a colorless crystalline solid (purity > 99.4%). CPF formulations in peanut oil (Millipore Sigma) were prepared twice weekly. Four CPF formulations (1, 3, 5, or 10 mg/kg) or an equal volume of the vehicle (peanut oil) were administered via subcutaneous (s.c.) injection for 21 consecutive days (study days 1-21), approximately 24 h apart. The injections were given along the back and the sites of administration were staggered to prevent local inflammation or built up of subcutaneous depots of oil. These exposures were carried out in their home cage. Rats were 9 weeks old at the initiation of dosing, and initial body weights ranged from 240-290 g. Animals were monitored daily for weight or discomfort throughout the study.

### ChE determinations

Systemically administered oily formulations of CPF are reported to form depot at the site of injection resulting in the slow release of the agent over the next couple of days [41]. Thus, to allow for this delayed pharmacokinetics, we waited for 48 h after the last CPF injection to assess the early effects of repeated CPF exposure on ChE levels. Following an 11-week washout period, these levels were reassessed in a separate cohort of rats. ChE determinations were made in the whole blood, hippocampus, and cortex. Briefly, rats were deeply anesthetized under isoflurane. Blood collected via cardiac puncture was transferred into chilled heparinized tubes and analyzed for ChE activity within 2 h of collection. Following blood collection, brains were rapidly dissected and the hippocampus and cortex were removed from each hemisphere, flash-frozen in liquid nitrogen, and stored at -80°C. ChE activity in whole blood, hippocampus, and cortex was determined following the standard Ellman reaction principle using a commercially available AChE activity assay kit (Catalog no. MAK119, Sigma-Millipore). The Ellman assay [42] was used to measure ChE activity in each brain region using 5,5’-dithio-bis (2-nitrobenzoic acid) (DTNB) as the colorimetric reagent and acetylthiocholine iodide (ASChI) as the AChE substrate. All samples were homogenized in lysis buffer (0.1 M phosphate buffer, pH 8.0 with 0.1% Triton) using a Polytron PT 1200 E and centrifuged for 1 min at 13,400 x g. The supernatant was collected and plated in triplicate in a 96-well plate for analysis. Blank wells contained an equal volume of buffer with DTNB. After equilibration with DTNB for 5 min, the reaction was started with the addition of ASChI. ASChI hydrolysis was quantified by measuring changes in absorbance at 405 nm over 15 min using the Synergy H1 Hybrid Plate Reader with Gen5 2.0 software (BioTek Instruments, Winooski VT, USA). ChE activity was normalized against total protein concentration determined using the BCA assay according to the manufacturer’s directions (Pierce, Rockford, IL, USA).

### Behavioral Assessments

To determine the long-term effects of repeated CPF exposures on neurobehavioral outcomes, CPF-treated rats along with age-matched vehicle-treated control rats were subjected to a battery of behavioral assays predictive of mood, anxiety, and depression-like condition. These assessments were carried out after an 11-week holding period post-CPF administrations. The behavioral tests included diverse assays such as the sucrose preference test (SPT), open-field test (OFT), elevated plus maze (EPM), and the forced swim test (FST). These assays were carried out in this same order thus going from the least stressful to the most stressful test. No two tests were carried out on the same day. Testing was carried out between 0800 to 1500 h and the testers were blind to treatment conditions.

#### Sucrose Preference Test (SPT)

This test measures hedonic (pleasure-seeking) or lack of it (anhedonia) by monitoring a rat’s preference for sucrose-laced water [43-46]. Briefly, rats were habituated to having two bottles in the cage lid for three days. The bottles were fitted with ball-bearing sipper tubes that prevented fluids from leaking. Following this acclimation, on the fourth day, rats were deprived of water but not food overnight (approx. 19-h). On the fifth day, two bottles were again introduced in the cage-top. Rats had the free choice of either drinking a 1% sucrose solution or plain water for a total period of 1-h. Both the volume of fluid and the weight of the bottle were measured before and after the test. Sucrose preference was calculated as a percentage of the sucrose intake over the total fluid intake.

#### Open Field Test (OFT)

This is a simple sensorimotor test used to determine a rat’s exploratory behavior, locomotor activity. By measuring rats’ preference to stay close to the walls of the field (thigmotaxis) versus time spent in the center of the arena, a gross assessment of animals’ anxiety could be made. Briefly, the arena consisted of a black Perspex box 90 × 60 × 50 cm kept in a dimly illuminated and quiet animal behavior testing room. A rat was allowed to explore the box uninterrupted for 10 min. The arena was cleaned with a 70% ethanol solution and dried completely in between each subject to eliminate any potential odor cues left by previous subjects. An overhead camera coupled to a computer running the animal tracking software (Noldus Ethovision XT 11) was used to record the test session. Using the software, a virtual grid was overlaid on the open field arena that divided the box into a central area and the surrounding area. The tracking software was used to measure time spent in the center along with other locomotor parameters such as the speed and distance covered in the arena by the animal.

#### Elevated Plus Maze (EPM)

This test assesses anxiety by considering the innate behavior of rats to prefer dark enclosed spaces over bright open spaces [43, 44, 47]. By using rodents’ proclivity toward dark, enclosed spaces (approach) and an unconditioned fear of heights/ open spaces (avoidance), EPM allow for the measurement of anxiety state in the absence of noxious stimuli that typically produce a conditioned response. The maze (Med Associates Inc., St. Albans, VT) was made of black polyvinyl chloride and consisted of four arms, 50 cm long x 10 cm wide, connected by a central square, 10 × 10 cm: two open without walls and two closed by 31-cm-high walls. All arms were attached to sturdy metal legs; the maze was elevated 55 cm above the floor level and was set in a dimly lit room. A video camera was suspended above the maze to record the rat movements for analysis. A video-tracking system (Noldus Ethovision XT 11) was used to automatically collect behavioral data. The procedure consisted of placing the rats at the junction of the open and closed arms, the center of the maze, facing the open arm opposite to where the experimenter was. The video-tracking system was started after the animal was placed in the maze so that the behavior of each animal was consistently recorded for 5 min. At the end of the 5 min test session, the rat was removed from the plus maze and returned to its home cage. The maze was cleaned with 70% ethanol and air-dried to remove any scent traces and allowed to dry completely before introducing the next animal in the arena. Time spent and entries made in the various arms of EPM were calculated.

#### Forced Swim Test (FST)

Porsolt’s modified FST was used to assess behavioral despair [43-45, 48]. Briefly, animals were placed in a glass cylindrical chamber (46cm H x 30cm D) filled with water (30 cm height, 25°C) forced to swim for 6-min. Swimming sessions were recorded for offline analysis. Immobility was defined as the period during which the animal floats in the water making only those movements necessary to keep its head above water was evaluated by two reviewers. The tank was emptied and thoroughly cleaned for every rat to be tested in a session.

### Data analysis

Data were analyzed, and graphs were plotted using the GraphPad software (Prism 9). Data were first examined for normality using Shapiro-Wilk’s tests. Variables not normally distributed were log-transformed and normality confirmed on the transformed data before analyses with *t*-tests. Normally distributed data were analyzed with independent t-tests or a One-way Analysis of variance (ANOVA) followed by a posthoc Tukey test wherever appropriate. In all cases, statistical significance was indicated by *p < 0.05.

## Results

### Dose-dependent effects of repeated CPF exposures on body weight

Daily administration of CPF at 1, 3, 5, or 10 mg/kg did not produce any overt toxicity or clinical signs of acute cholinergic crisis during the 21-day exposure period. No significant differences in the body weight gain were observed between vehicle controls and various CPF groups until day 20 when the bodyweight in the CPF-10 group was significantly less on day 21 compared to the vehicle group (*p* = 0.04, RM-ANOVA, Fig. 1). This minor weight difference was transient and measurements after the exposure period did not reveal any significant differences in the body weight between vehicle and CPF groups.

**Figure 1.**
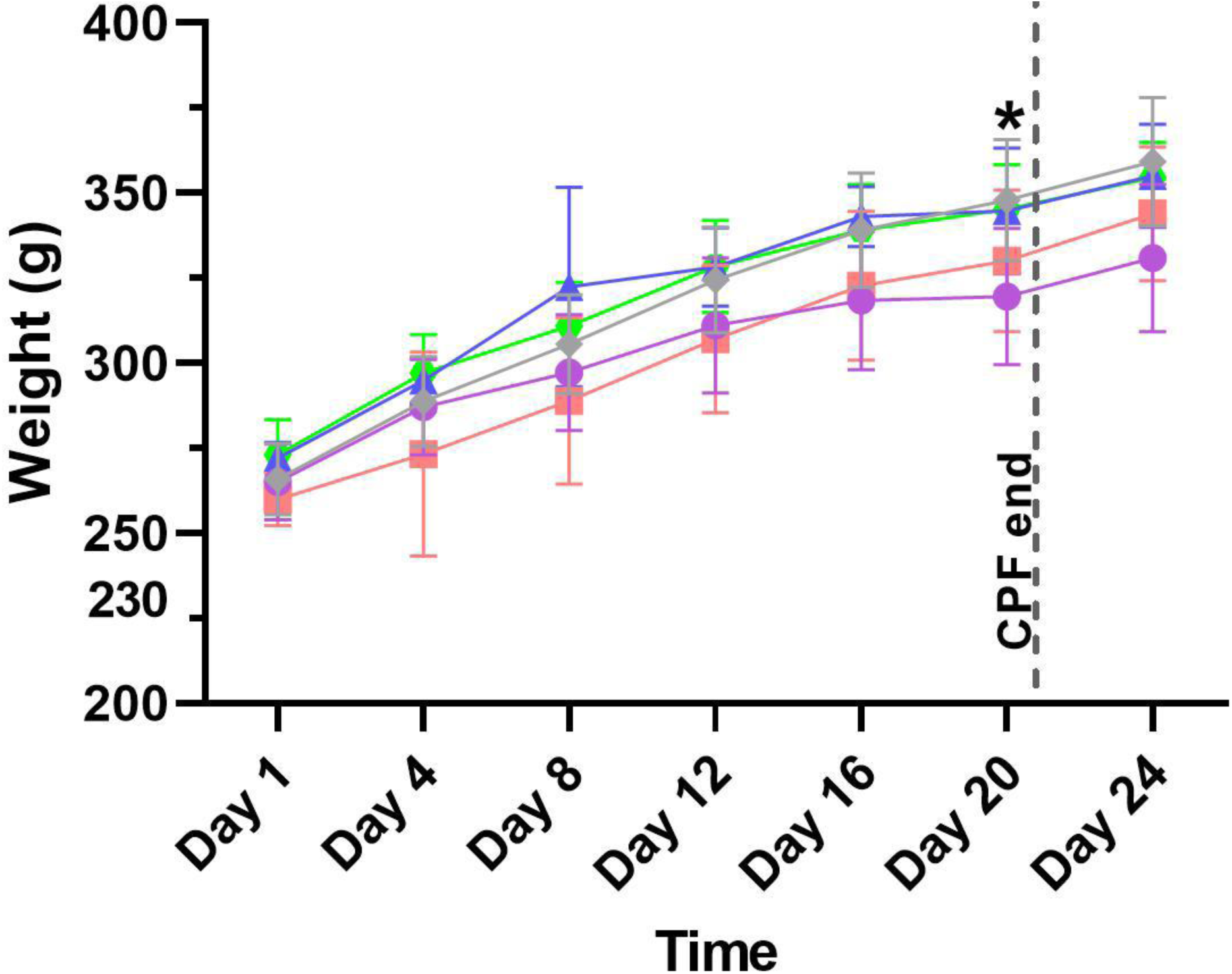
Effects of repeated CPF exposures on body weight. Bodyweight measurements for Sprague-Dawley rats treated with vehicle or various doses of CPF. No significant differences in the weights were noted across all the CPF groups compared to vehicle-treated rats till day 20 of the CPF exposures. On Day 21, statistically significant differences between vehicle-treated and CPF-10 treated groups were noted. Weight measurements on the following days did not exhibit continual significant differences (Data expressed as mean ± S.D., n= 12 rats/ group, *p<0.05, repeat measures ANOVA) [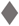- Vehicle, 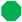- CPF1, 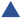- CPF3, 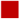- CPF5, 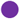- CPF10].

### Dose-dependent effect of repeated CPF exposures on ChE activity

Blood and brain ChE activity was assessed at 2 days after the end of the 21-day CPF exposure paradigm. ChE activity was significantly lower in whole blood, hippocampus, and cortex in all of the CPF-exposed groups relative to control. As shown in Fig 2, regression analyses revealed a strong correlation between CPF doses and ChE activity in blood (r^2^= 0.88, p= 0.05), cortex (r^2^= 0.73, p= 0.1) and hippocampus (r^2^= 0.89, p= 0.03) respectively. Thus, blood ChE activity levels were approximately 36%, 26%, 22%, and 15% of vehicle control levels following administration of CPF at 1, 3, 5, or 10 mg/kg/d respectively. ChE activity levels in the cortex were approximately 98%, 55%, 54%, and 38% of vehicle controls while, hippocampal ChE activity levels were approximately 78%, 75%, 60%, and 51% of vehicle controls following administration of CPF at 1, 3, 5 or 10 mg/kg/d respectively.

**Figure 2.**
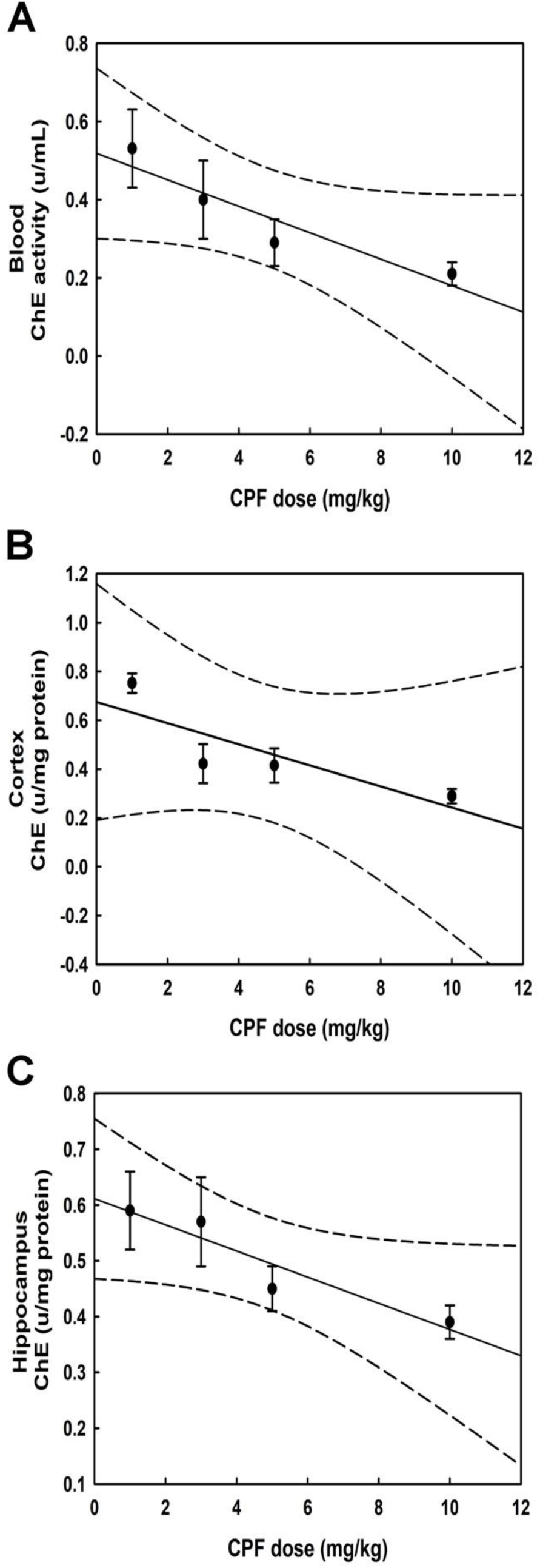
Blood and brain Cholinesterase (ChE) activity level early point following repeated CPF exposures. CPF was administered daily for 21 days at 1, 3, 5, and 10 mg/kg doses respectively and ChE activity was assessed early point (2-day post) following the last CPF dose. Correlation analysis revealed a significant association between CPF dose and ChE activity in A. blood (r^2^= 0.88, p= 0.05), B. cortex (r^2^= 0.73, p= 0.1) and C. hippocampus (r^2^= 0.89, p= 0.03) respectively at 2-days after the last CPF dose.

ChE activity levels, when reassessed following an 11-week washout period (Fig. 3) after the end of the 21-day CPF exposure paradigm, indicated no significant depression in ChE activities in the blood (p= 0.25) or cortex (p= 0.55) or hippocampus (p= 0.14) for any of the CPF groups relative to vehicle control (Group data are shown with boxes representing the 25–75 percentile; horizontal lines in the boxes represents the median; and whiskers indicate the maximum and minimum values (n= 4-5 rats/ group, *p<0.05, One-way ANOVA).

**Figure 3.**
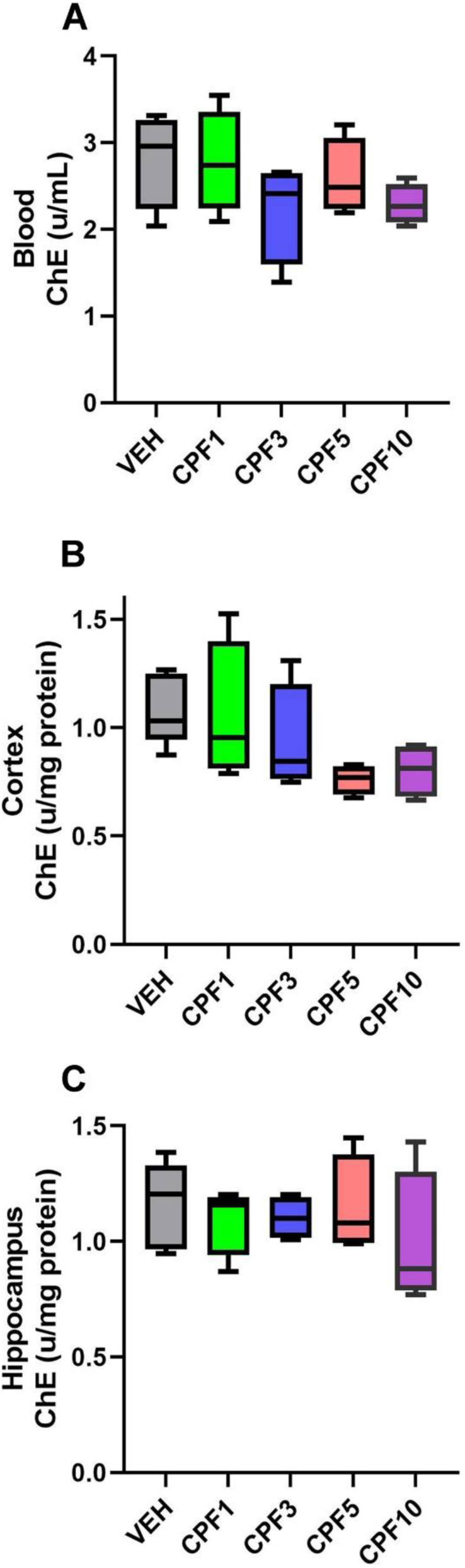
Blood and brain Cholinesterase (ChE) activity level at a late time point following repeated CPF exposures. CPF was administered daily for 21 days at 1, 3, 5, and 10 mg/kg doses respectively and ChE activity was assessed in the blood and the brain 11 weeks (late time point) following the last CPF dose. No significant differences in ChE activity were found amongst the experimental groups in the A. blood, B. cortex, and C. hippocampus (Group data are shown with boxes representing the 25–75 percentile; horizontal lines in the boxes represents the median; and whiskers indicate the maximum and minimum values (n= 4-5 rats/ group, *p<0.05, One-way ANOVA).

### Dose-dependent effects of repeated CPF exposures on SPT performance

The percent fluid consumptions were calculated for the vehicle-treated and the CPF groups. Data revealed a significant preference for sucrose-laced compared to normal water among vehicle (72.8 ± 7.8), CPF-1 (68.4 ± 13.6), and CPF-3 (69.9 ± 9.3) groups whereas, CPF-5 (55.1 ± 23.3) and CPF-10 (49.7 ± 7.7) groups did not exhibit significant preference towards sucrose indicative of an anhedonia-like state (*p<0.05, t-test, n=6 rats/group, mean ± S.D., Fig. 4A). We next compared percent sucrose preference across various groups (Fig. 4B). A One-way ANOVA revealed significant differences in sucrose intake between the groups. Post-hoc analysis showed that CPF-5 and CPF-10 treated rats exhibited significantly lower sucrose preference compared to the vehicle-treated group (*p<0.05, One-way ANOVA, Dunnett’s posthoc test, n= 6 rats/group, mean ± S.D., Fig. 4B). No significant differences (p= 0.2, Fig. 4C) were observed between total fluid consumption between the experimental groups (Group data are shown with boxes representing the 25–75 percentile; horizontal lines in the boxes represents the median, and whiskers indicate the maximum and minimum values; n= 6 rats/ group, *p<0.05, One-way ANOVA, Dunnett’s posthoc test).

**Figure 4.**
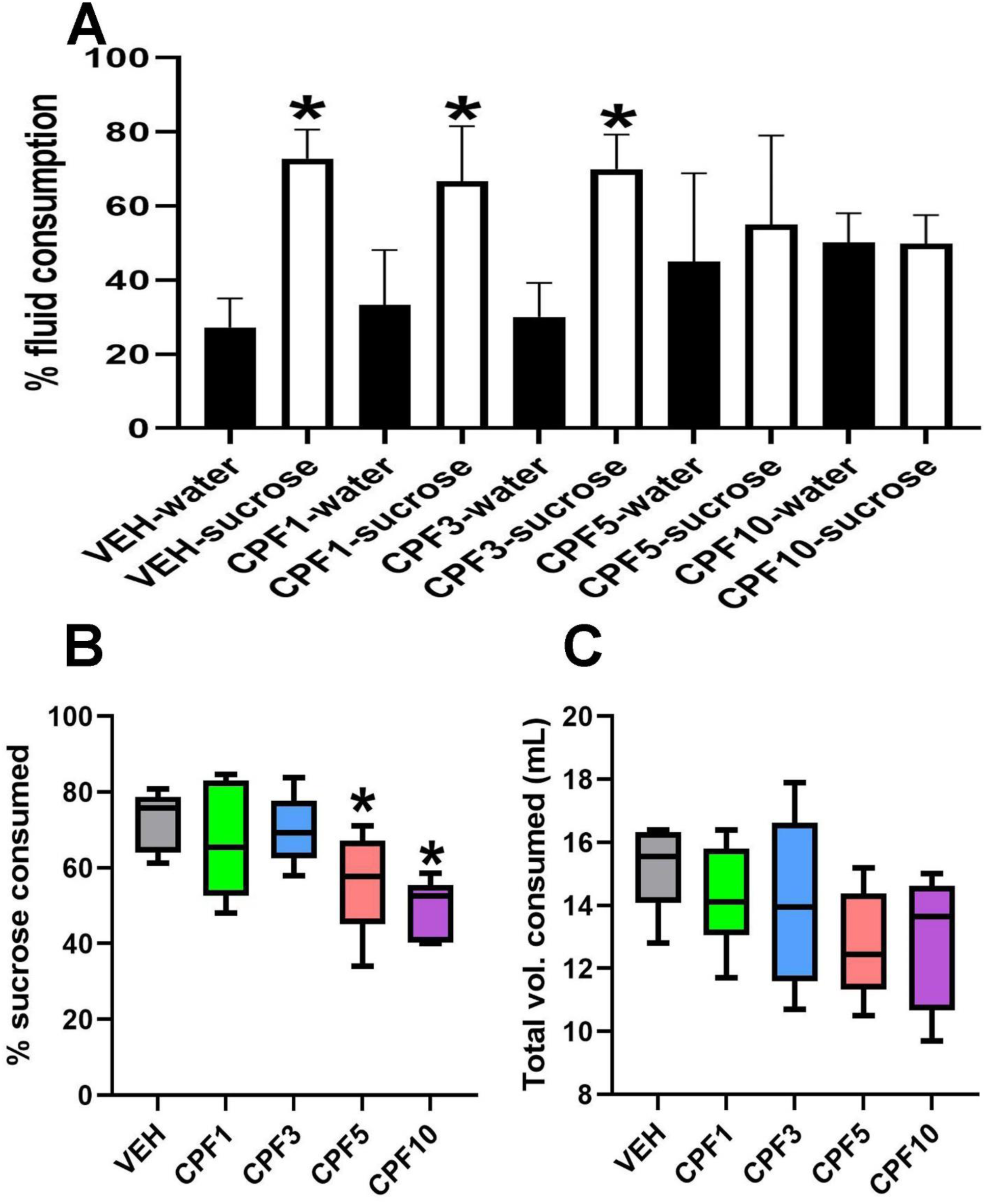
Effect of repeated CPF exposures on Sucrose preference. **A**. Bar graph depicting % fluid consumed (water vs sucrose) within in each group. Rats in the vehicle, CPF-1, and CPF-3 groups demonstrated a significantly greater preference for sucrose consumption over regular water whereas rats in the CPF-5 and CPF-10 did not exhibit any such preference indicating an anhedonia-like condition (Data expressed as mean ± S.D., n= 6 rats/ group, *p<0.05, t-test comparing water vs sucrose within each group). **B**. Analyses of % sucrose consumed across all the experimental groups revealed that rats in the CPF-5 and CPF-10 group exhibited significantly lower sucrose consumption compared to the vehicle-treated rats. **C**. The total volume consumed (water + sucrose) was not significantly different between the groups (Group data are shown with boxes representing the 25–75 percentile; horizontal lines in the boxes represents the median; and whiskers indicate the maximum and minimum values; n= 6 rats/ group, *p<0.05, One-way ANOVA, Dunnett’s posthoc test).

### Dose-dependent effects of repeated CPF exposures on OFT performance

Fig. 5 illustrates the chronic effect of repeated CPF exposures on OF locomotor activity. A dose-dependent decrease in the time spent in the center (Fig. 5B) along with a decrease in the number of visits to the center (Fig. 5C) were noted in CPF-treated rats. A One-way ANOVA revealed significant differences in OF performance amongst the various tested groups. Post-hoc analyses revealed that CPF 1 and 3 mg/kg groups were not significantly different from the vehicle-treated rats while CPF-5 and 10 mg/kg rats demonstrated significant decreases in time-spent and visits to the center of arena compared to vehicle-treated rats. No significant differences in distance moved (p= 0.5, Fig. 5D), and speed (p= 0.2, Fig. 5E) were observed between the various groups suggesting no involvement of motor deficits in the OFT outcomes (Group data are shown with boxes representing the 25–75 percentile; horizontal lines in the boxes represents the median, and whiskers indicate the maximum and minimum values; n= 6 rats/ group, *p<0.05, One-way ANOVA, Dunnett’s posthoc test).

**Figure 5.**
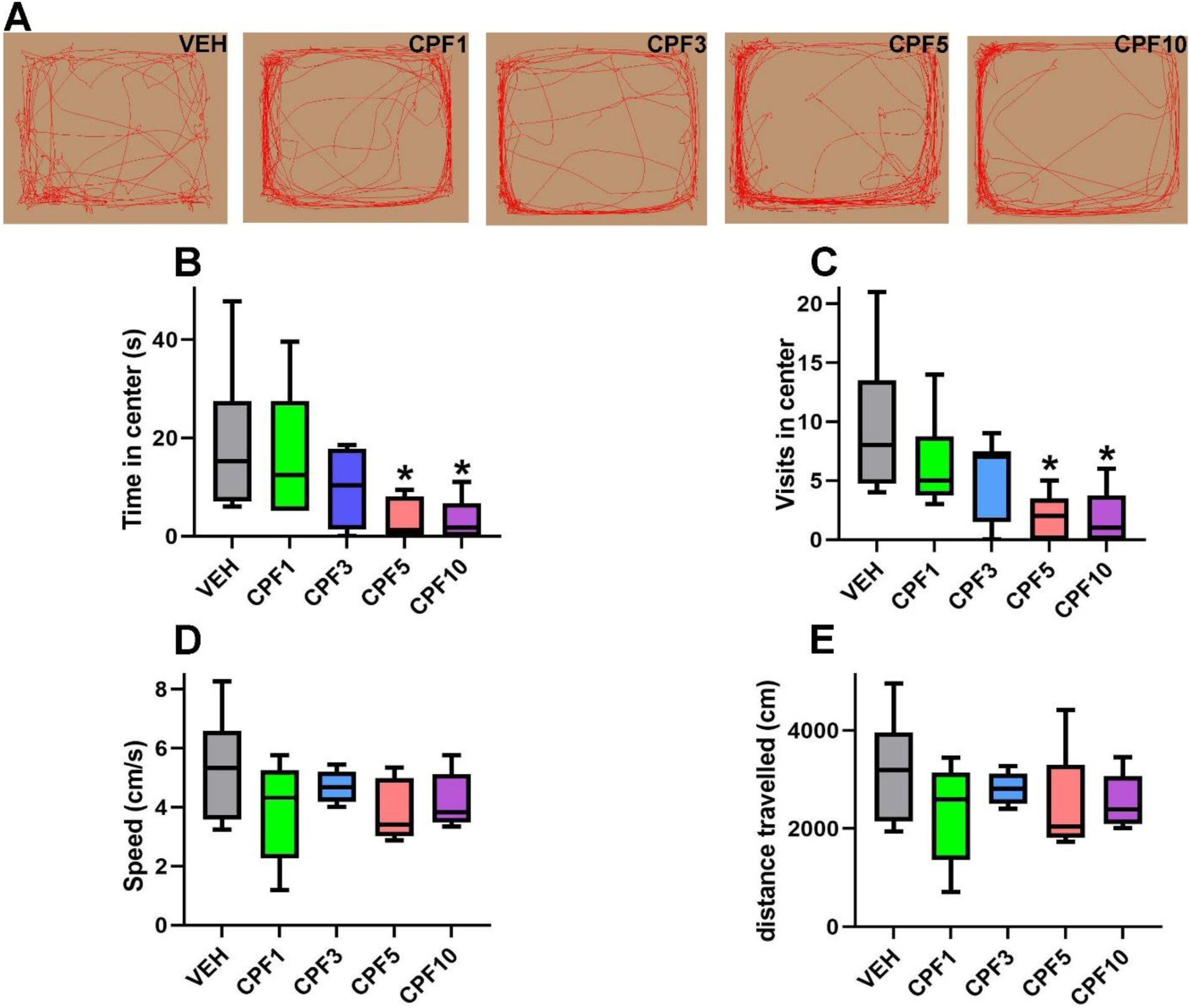
Effect of repeated CPF exposures on Open-field performance. **A**. Representative activity profile of rats treated with vehicle (VEH) and various CPF doses on the Open-field exploration. With increasing doses of CPF, a decrease in a visit to the center of the arena and a preference to stay along the boundaries of the arena could be noted. A significant reduction in time spent in the center of the arena (B) along with a corresponding decrease in the number of visits in the center of the arena (C) was noted in rats belonging to CPF-5 and CPF-10 groups compared to vehicle-treated rats. **D. and E**. No significant changes between speed and distance moved in the open-field arena were noted between the VEH-treated and various doses of CFP-treated rats (Group data are shown with boxes representing the 25–75 percentile; horizontal lines in the boxes represents the median, and whiskers indicate the maximum and minimum values; n= 6 rats/ group, *p<0.05, One-way ANOVA, Dunnett’s posthoc test).

### Dose-dependent effects of repeated CPF exposures on EPM performance

Comparing the time-spent and entries made in the open and closed arm of the EPM allows for the assessment of anxiety-like state in rodents. Increased close-arm exploration is considered to indicate the anxiogenic state in rats. Data presented in Fig. 6 shows that CPF dose-dependently produced decreases in open-arm time (Fig. 6B) and an increase in close-arm time (Fig. 6C) on the EPM. A One-way ANOVA revealed significant differences amongst the various tested groups. Post-hoc analyses revealed that CPF 1, 3, or 5 mg/kg groups were not significantly different from the vehicle-treated rats while CPF-10 rats demonstrated significant increases in close-arm time and corresponding decreases in open-arm time compared to vehicle-treated rats. Further, we observed a dose-dependent trend towards a decrease in open-arm entries (Fig. 6D) and an increase in close-arm entries (Fig. 6E). But, statistical significance was not noted. Finally, no significant differences in distance moved (p= 0.4, Fig. 6F), and speed (p= 0.5, Fig.6G) were observed between the various groups suggesting no involvement of motor deficits in the EPM outcomes (Group data are shown with boxes representing the 25–75 percentile; horizontal lines in the boxes represents the median, and whiskers indicate the maximum and minimum values; n= 6 rats/ group, *p<0.05, One-way ANOVA, Dunnett’s posthoc test).

**Figure 6.**
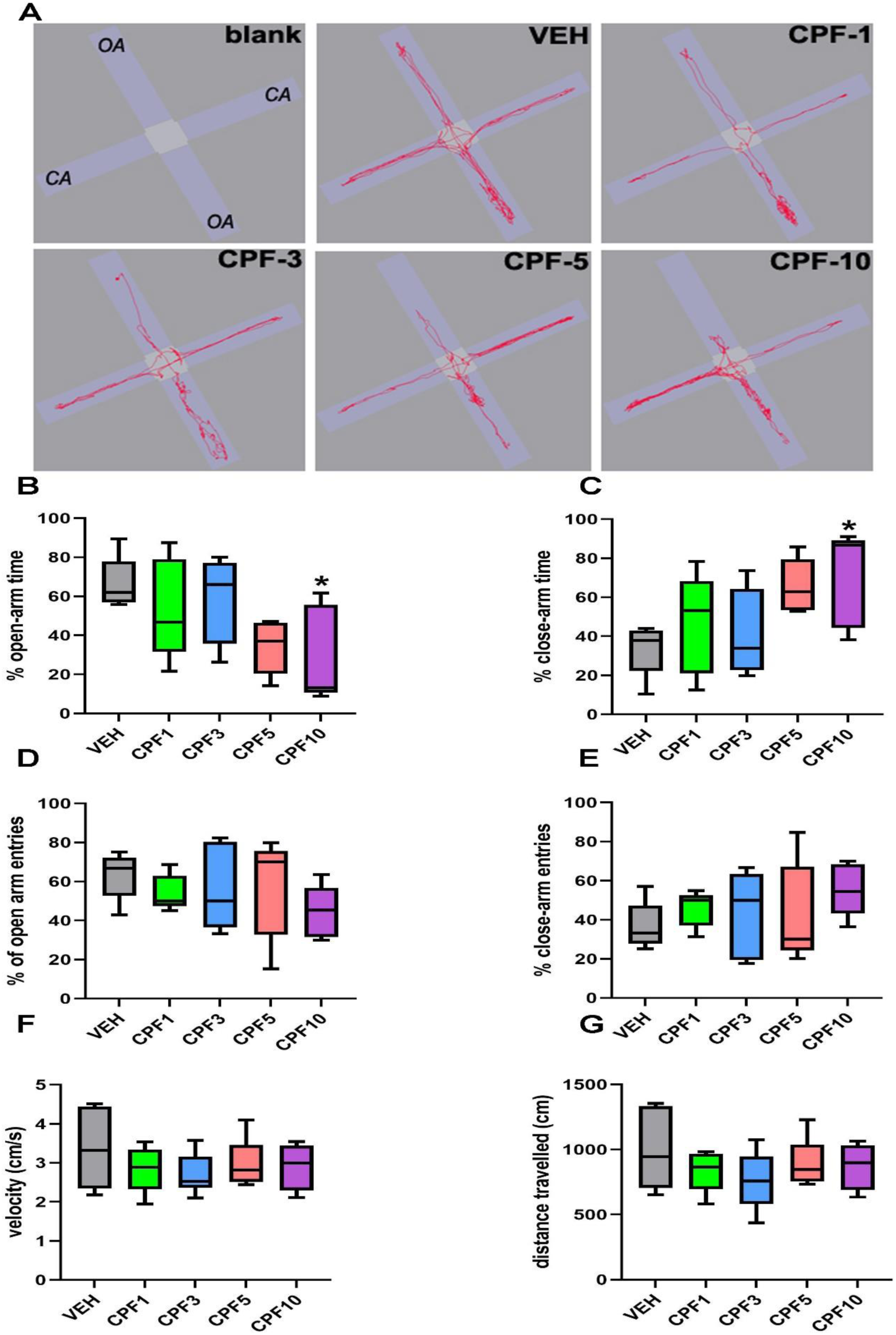
Effect of repeated CPF exposures on EPM performance. **A**. Representative activity profile of rats treated with vehicle (VEH) and various CPF doses on the EPM. Blank shows the orientation of the Plus Maze with open arms represented as OA and closed arms represented as CA. **B. and C**. A dose-dependent reduction in % time-spent in the OA and an increase in % time-spent in the CA was noted in CPF groups but a significantly different OA vs CA preference was noted only for the CPF-10 treated rats. **D. and E**. While the % OA entries decreased and a corresponding increase in % CA entries was noted for the CPF-10 treated rats, these data were not significantly different from the VEH-treated rats. **F. and G**. No significant changes between velocity and distance moved on the EPM were noted between the VEH-treated and various doses of CFP-treated rats (Group data are shown with boxes representing the 25–75 percentile; horizontal lines in the boxes represents the median; and whiskers indicate the maximum and minimum values; n= 6 rats/ group, *p<0.05, One-way ANOVA, Dunnett’s posthoc test).

### Dose-dependent effects of repeated CPF exposures on FST performance

Despair is a common sign of depression and FST performance is a widely used measure of a despair-like state in rodents. Vehicle-treated rats exhibited an immobility time of 110.6 ± 19.8 s. CPF groups exhibited a dose-dependent increase in immobility time. Thus, immobility times were CPF-1 (137.9 ± 25.3 s), CPF-3 (178.1 ± 36.9 s), CPF-5 (235.0 ± 18.4 s) and CPF-10 (227.9 ± 18.0 s) respectively. One-way ANOVA revealed significant differences amongst the groups. Post-hoc Dunnett test revealed that CPF 3, 5, and 10 mg/kg rats exhibited significantly greater immobility time indicative of signs of despair compared to age-matched, vehicle-treated rats (Group data are shown with boxes representing the 25–75 percentile; horizontal lines in the boxes represents the median, and whiskers indicate the maximum and minimum values; n= 6 rats/ group, *p<0.05, One-way ANOVA, Dunnett’s posthoc test, Fig. 7).

**Figure 7.**
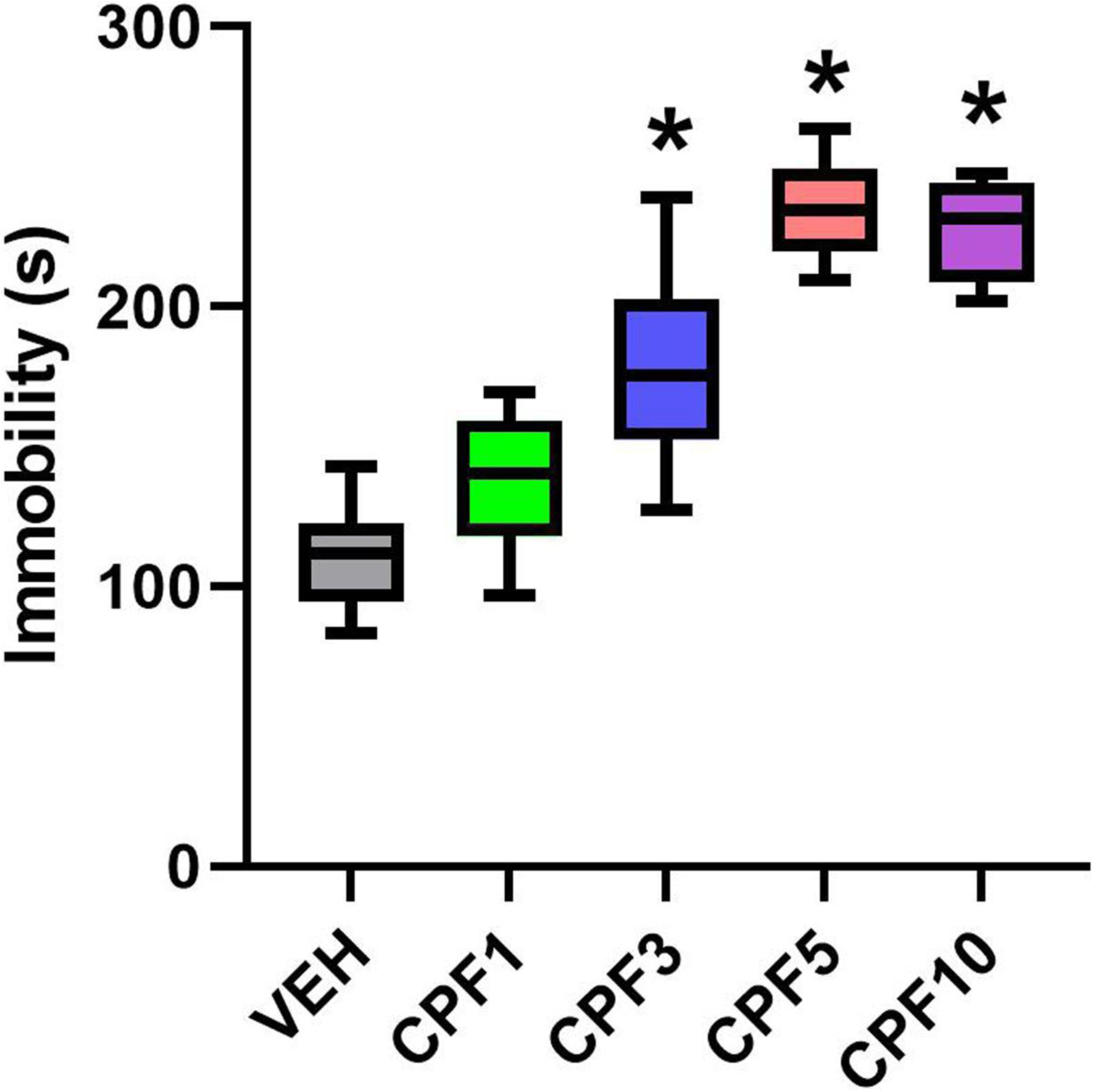
Effect of repeated CPF exposures on FST performance. A dose-dependent increase in signs of behavioral despair was noted in CPF treated rats. A significantly greater immobility time was noted in rats repeatedly dosed with 3, 5, and 10 mg/kg/d of CPF compared to vehicle-treated rats (Group data are shown with boxes representing the 25–75 percentile; horizontal lines in the boxes represents the median, and whiskers indicate the maximum and minimum values; n= 6 rats/ group, *p<0.05, One-way ANOVA, Dunnett’s posthoc test).

## Discussion

To provide an experimental paradigm for assessing latent effects of occupational OP exposures on the development of a depression-like condition, we systematically planned a study with the following criteria. First, we used CPF as our prototype OP pesticide. CPF is amongst one of the most widely used pesticides around the world in occupational, commercial, and residential settings [16, 29]. Thus, CPF is an environmentally relevant pesticide and a strong candidate to study the effects of OP exposures on biological systems. Second, we used adult rats for OP exposures since that age would be more relevant to populations that would more likely experience occupational-like exposures. Moreover, many elegant studies had already tested the effects of gestational or developmental, or adolescent OP exposures in rodents [21, 32-36, 49]. Third, we utilized a subcutaneous route for CPF exposures. Dermal exposure is a dominant route for OP exposures. Furthermore, subcutaneous OP exposure provides for a greater degree of dose control allowing for a reproducible pharmacological response [41, 43, 50] compared to other routes of administration such as oral [41] or inhalational [51]. Fourth, we used low CPF doses (1 mg/kg/d to 10 mg/kg/d) that were not associated with acute hypercholinergic response and thus would be considered sub-threshold doses [52]. Fifth, for the duration of OP exposure we chose a once-daily CPF injection protocol that continued for 21 consecutive days. This dosing paradigm in rats [50] has been shown to produce ChE inhibitions that mimic ChE activity levels seen following sub-chronic OP exposures in humans [53]. Finally, to study the latent effects of these repeated low-dose CPF exposures, rats were tested for signs of depression and anxiety at 11-weeks post-exposure. Our approach of combining repeated sub-threshold doses coupled with long-term behavioral assessment provides an experimental paradigm for studying cause and effect relationships between environmental and occupational OP exposures and the development of chronic behavioral deficits.

Previous studies have reported acute reductions in ChE activity upon repeated OP exposures, followed by the emergence of latent behavioral deficits despite the recovery of ChE activity to normal levels. The degree and time course of ChE inhibition and recovery observed in our study was comparable to others in this area. For example, CPF (18 mg/kg/d) administered on alternate days for 30 days to adult rats produced about 80% inhibition in plasma and approximately 50% inhibition in brain ChE activity at the end of the treatment period which, recovered to normal levels over the subsequent 7-week period [40]. In our study, the highest dose for CPF (10 mg/kg/d) produced up to ∼60% inhibition in brain ChE activity and up to ∼85% inhibition in blood ChE activity when assessed two days after the 21-day exposure paradigm. The comparable profiles of blood and brain ChE inhibition and recovery in these studies indicate that blood ChE activity can serve as a biomarker of brain ChE activity. Overall, the levels of ChE inhibition seen in our study agree with reported CPF doses that are considered sub-threshold and do not produce overt cholinergic response [52], and are in line with those reported in an occupational OP exposure setting [53].

In our study, CPF doses that were associated with a significant depressive phenotype (5 and 10 mg/kg/d) showed hippocampal ChE activity inhibited by approximately 40% and 50% respectively at the early time point, while cortical ChE activity at the early time point was inhibited by approximately 45% and 60% respectively. CPF doses that were not associated with the depressive phenotype (1 mg/kg/d) showed hippocampal and cortical ChE activity inhibited by approximately 20% and 2% respectively. The CPF 3 mg/kg dose, which elicited signs of depression in some behavioral tests, showed hippocampal and cortical ChE activity inhibited by 25% and 40% respectively at the early time-point. This suggests that signs of depression and memory impairment require some degree of initial ChE inhibition. However, given that these behavioral deficits persist beyond the point that ChE levels return to normal, other mechanisms beyond cholinergic stimulation must be at play to sustain these deficits.

Several authors have speculated that both cholinergic and non-cholinergic mechanisms could play a role in the development of chronic OP-related behavioral deficits [52, 54-58]. Among these mechanisms, neuroinflammation has been repeatedly demonstrated as an important mechanism for chronic OP neurotoxicity following high-dose OPs [59]. For example, repeated exposures to OPs such as CPF [60] and malathion [61] was reported to upregulate the hippocampal expression of the glial fibrillary acidic protein (GFAP) in mice while, microglial activation and increases in proinflammatory cytokines have been reported following dichlorvos administration in rats [62, 63]. Clinical and laboratory studies have indicated chronic neuroimmune dysfunctions as key mediators of depression symptoms and memory difficulties in GWI [12, 14]. Similarly, repeated exposure to OP compounds including CPF and DFP produce chronic alterations in components of neurotransmission [64] including second-messenger calcium elevations [65, 66] and impaired axonal transport [67-69]. Chronic upregulation of glutamatergic, dopaminergic, and serotonergic genes has been reported following repeated OP diazinon exposures in rats [70]. Furthermore, alterations in the hippocampal expression of neuropeptides and neurotrophic factors including the brain-derived neurotrophic factor (BDNF) have been reported following repeated low-dose CPF [71, 72] or DFP exposures [73]. This is particularly interesting because BDNF is both implicated in depression pathophysiology and antidepressant response [74, 75]. Moreover, outside of its trophic role, the role of BDNF in affecting synaptic plasticity and neurogenesis is critical [74, 76]. It is therefore important to note that repeated CPF (5 mg/kg/d for 5 days) was associated with a chronic decreased dendritic spine densities and synaptic plasticity in mice [77]. Similarly, decreased adult neurogenesis has been reported following exposure to various pesticides and agrochemicals [78, 79]. It will be interesting to study the latent effects of sub-chronic, sub-threshold CPF on synaptic plasticity, neurogenesis and development of depression.

In recent times, genetic [80] and epigenetic [81] mechanisms have been highlighted as potential mediators of OP-induced chronic neurotoxicity. For example, individuals with low activity variants of the Paraoxonase (*PON1*) gene that hydrolyzes active metabolites of OPs may be more susceptible to ill-effects of OP exposure [82, 83]. Epigenetic mechanisms may mediate the long-term consequences of OP exposures [84]. For example, in humans, prenatal CPF exposure measured via cord blood was associated with altered DNA methylation levels at the PPARγ gene, in addition to poorer performance in cognitive and language domains at age two [85]. Ambient OP exposure, as assessed via residential and workplace proximity to commercial applications, was associated with differential DNA methylation at several loci in the genome of older adults, with muscarinic acetylcholine receptor pathways being among the most affected [86]. In model organisms, several studies have shown epigenetic consequences of exposure to CPF [49, 87-89] and OP compounds more generally [81]. Considering OP-induced epigenetic and behavioral changes, we previously showed that chronic, low-dose DFP exposure led to upregulation of histone deacetylases and decreased levels of histone 3 lysine 9 acetylations (H3K9ac) at *Bdnf* promoter IV in the rat hippocampus, possibly contributing towards chronic reductions in BDNF protein levels following such low-dose OP exposures. This may be one mechanism through which OP affects mood and depressive outcomes. Overall, these studies in diverse tissues (e.g. blood, brain) and organisms show epigenetic consequences of OP exposure. However, the mechanisms through which OP compounds elicit epigenetic changes remain to be fully elucidated.

In summary, we have demonstrated the persistence of depressive signs following repeated low dose CPF exposures in adult rats. This could result from myriad molecular mechanisms such as changes to second messenger systems, gene expression, epigenetic modifications, or synaptic plasticity. The CPF exposure paradigm proposed in this study could provide an ideal model system to further study these molecular mechanisms.

